# Levenshtein Distance as a Measure of Accuracy and Precision in Forensic PCR-MPS Methods

**DOI:** 10.1101/2021.01.03.425149

**Authors:** Brian Young, Tom Faris, Luigi Armogida

**Author notes:** Corresponding Author: Dr. Brian Young, NicheVision Forensics, LLC, 526 South Main Street, Akron, Ohio, 44311.

## Abstract

Accuracy and precision determinations are standard components of method validations where they help to describe the performance of methods. Despite their importance, a standard approach to calculating these parameters is not available for forensic PCR-MPS methods that detect sequence-based alleles. In this paper, we describe a method based on the Levenshtein distance metric which aptly summarizes method accuracy in terms of the closeness of read sequences to reference sequences, and method precision in terms of the agreement among read sequences. Inaccuracy or imprecision in forensic methods can lead to wrong allele calls. By expressing method performance in terms of a distance metric, this method places PCR-MPS on equal footing with distance-based measures in PCR-CE methods. Summary statistics based on the Levenshtein distance can be used to compare performance of different kits, markers, sequencers, or methods.

## 1. Introduction

Performance evaluations of quantitative analytical methods used in forensics often includes consideration of accuracy and precision [1]. These measures are particularly useful when describing the suitability of a method in an intended application or when comparing the performance of alternative methods [2]. Method accuracy and precision are core components of forensic method validations under SWGDAM and ENFSI guidelines [3–5]. In this context, accuracy has been defined as the degree of conformity of measurements to the true value; and precision has been defined as the degree of mutual agreement among a series of individual measurements [3].

### 1.1. Accuracy and Precision in PCR-CE Methods

The goal of PCR-CE methods is to determine the true length of PCR-targeted markers in the genome. In these methods, the measurements of interest are electrophoretic retention times from which the lengths of PCR amplicons can be derived. Electrophoretic retention times are routinely transformed into fractional base pair units. Even though “base pair” is a discontinuous measure, the fractional units are essentially continuous. Means and standard deviations are calculated in units of fractional base pair usually under an assumption that Normal distributions fit the data [6, 7]. To avoid length-miscalls, PCR-CE methods must be accurate to retention times that are within ± 0.5 base pairs of the true amplicon length [5, 8]. It is not possible to measure accuracy at the individual PCR amplicon level. Rather, the entity measured is the mode of retention times of an aggregate of same-length amplicons that migrate together under electrophoresis [9]. Precision is commonly described in terms of a standard deviation of measurement error of the aggregate [10, 11]. That is, the standard deviation of replicate electrophoretic peak retention times.

In addition to error present in the electrophoretic process, PCR-CE methods can introduce error in PCR amplicons due to PCR misincorporation and PCR stutter. PCR misincorporation possibly contributes to variation in electrophoretic retention times. Stutter error holds a special status in forensic STR analysis. Stutter peaks are expected in electropherograms, and presence (or absence) of stutter contributes to visual assessments of the sensitivity of the method. Stutter artifacts are not typically considered in measures of accuracy or precision in PCR-CE methods.

### 1.2. Accuracy and Precision in PCR-MPS Methods

In contrast to PCR-CE methods, the goal of PCR-MPS methods is to determine the nucleotide sequence of PCR-targeted markers in the genome. Experimental determination of a marker sequence results in a sequencer read (hereafter simply called a read) expressed as a string of letters following the IUPAC coding convention [12] for the four nucleotides in the DNA backbone (A, C, G, T)^1^. Like PCR-CE, the genomic segments targeted by the assay are defined by the PCR primers, which for most STR markers, includes flanking sequence that may contain additional polymorphisms. Targeted segments can be bioinformatically trimmed to shorter extents, and typically at least the PCR primer sequences are trimmed away. The read sequence that remains becomes the sequence-based allele under consideration. Like PCR-CE, accuracy and precision should be sensitive to single nucleotide differences^2^ because authentic allele sequences can differ by a single nucleotide due to point polymorphisms [13, 14]. PCR-MPS methods generate a read sequence for each individual PCR amplicon, and a base call for each nucleotide position. Thus, accuracy and precision can be determined at several levels: nucleotide, read, and allele.

Consensus has not yet been reached for reporting accuracy and precision for forensic PCR-MPS methods. Accuracy of PCR-MPS methods has been reported either as a call rate [15] or as a concordance to CE allele numbers. Precision is often not reported at all [16–21] or has been reported as a modified call rate [15].

### 1.3. Distance Measures

Accuracy and precision are often expressed in distance units. The measurement result of a PCR-MPS method is a read string which may be close to or distant from the expected sequence. The distance between two DNA sequences is called an edit distance. The concept of edit distance is tightly associated with sequence alignment, which can be thought of as a transcript of the edits (nucleotide substitutions, insertions, and deletions) required to convert one sequence string into another [22] (p 125). An optimized alignment may not correspond to the minimum possible number of edits due to weightings that may be assigned to mismatches and gaps in the aligned sequences. The minimum number of equally-weighted edit operations is called the Levenshtein distance [22, 23].

Levenshtein distance has been shown to satisfy [24] the axioms [25] (p.147) required of distance metrics (Appendix A). Distance metrics imply the existence of an underlying metric space. The concept of a metric space where the elements of the space are DNA (or protein) sequence strings has a well-established foundation. Maynard Smith [26] first described sequence spaces in the context of natural selection, while Eigen [27] provided a formal topological/geometrical interpretation of sequence spaces. In short, Levenshtein distance is a distance metric that can be used in mathematical and statistical calculations. By employing the Levenshtein distance metric, accuracy and precision can be expressed in accord with SWGDAM guidance. That is, accuracy in amplicon sequencing can be expressed as the degree of conformity of read sequences to the true DNA sequence, and precision can be expressed as the degree of mutual agreement among a series of individual read sequence determinations.

Here, we define concepts for accuracy and precision in sequence-based PCR-MPS methods that are 1) compliant with SWGDAM and ENFSI guidelines, 2) analogous to the concepts of sizing accuracy and precision currently used for PCR-CE methods [28], and 3) can be expressed in units that can be used in statistical calculations.

## 2. Material and Methods

### 2.1. Samples and Sequencing

One ng aliquots of four DNA standards were sequenced in triplicate: 2800M (Promega, Madison, WI), and NIST standard reference material 2391d components A, B, and E (NIST, Gaithersburg, MD). Sequencing was performed at Intermountain Forensics (Salt Lake City, UT) using the ForenSeq™ kit and MiSeq™ sequencer (Verogen, San Diego, CA). Raw FASTQ data files were analyzed using MixtureAce™ (NicheVision Forensics, Akron, OH) with an analytical threshold of zero. Read sequences were bioinformatically trimmed to match the Verogen recommended analyzable regions [29] and exported to comma separated value (CSV) files along with their read count intensities (RCI). Stutter artifacts identified by MixtureAce were deleted from the CSV files. A custom Python script was used to modify the CSV files for statistical analysis such that each data row corresponded to an individual sequencer read (RCI = 1). Sixteen alleles were selected for analysis based on being homozygous in one of the DNA standards (Table 1). Selecting homozygous alleles allowed unambiguous association of erroneous sequences to parent alleles.

**Table 1.** Alleles selected for analysis.

| Standard<sup>1</sup> | Locus | Allele ID<sup>1</sup> | STRSEQ<sup>2</sup><br>Accession No. |
| --- | --- | --- | --- |
| 2800M | CSF1PO | 12 PD | MH085189.1 |
|  | D5S818 | 12 RR | MH167007.1 |
|  | D22S1045 | 16 AW | MH167279.1 |
|  | TPOX | 11 UN | MF044255.1 |
| NIST SRM 2391d<br>Component A | D2S441 | 11 RC | MH167320.1 |
|  | D3S1358 | 17 ZU | MH166980.1 |
|  | D4S2408 | 9 ZI | MH039589.1 |
| NIST SRM 2391d<br>Component B | D7S820 | 10 TO | MH167034.1 |
|  | D13S317 | 11 AQ | MH167219.1 |
|  | TH01 | 7 CS | MH085116.1 |
| NIST SRM 2391d<br>Component E | D6S1043 | 11 BZ | MH166925.1 |
|  | D9S1122 | 11 KZ | MH085132.1 |
|  | D10S1248 | 14 TJ | MH167062.1 |
|  | D19S433 | 14 WU | MH174820.1 |
|  | D20S482 | 15 QX | MH085169.1 |
|  | PentaD | 14 WB | MH232690.1 |

<sup>1</sup> Allele identifiers consist of the CE-equivalent allele number and the first two digits of the SID label [30].
<sup>2</sup> https://www.ncbi.nlm.nih.gov/bioproject/380127

Reads exhibiting Levenshtein distances > 10 were omitted from the data set. Reads with extreme Levenshtein distances may arise from trace level contamination, or PCR mis-priming of non-human sequence, and therefore are not representative data. Of 125,994 reads in the dataset, only 107 exhibited Levenshtein distances of > 10 from the expected sequences. The maximum Levenshtein distance was 78. The truncated data set included 125,887 reads. Levenshtein distances were calculated using a custom Python script employing the Jellyfish module (https://github.com/jamesturk/jellyfish). All other data analysis was performed using custom R scripts.

## 3. Results

### 3.1. Distribution of Levenshtein Distances

Histograms of Levenshtein distances of the individual reads generated by a PCR-MPS method exhibit a characteristic shape independent of locus (Figures 1-2; Appendix B). Most reads match the expected sequence and therefore, exhibit a Levenshtein distance of zero from the expected sequence. The most abundant error type consists of reads exhibiting a single point error and a corresponding Levenshtein distance from expected equal to one. Read frequencies tend to decline exponentially with increasing Levenshtein distance (Figures 1-2). If point errors in reads arise randomly, read count distributions would be expected to be Poisson. However, it is known that sequencing error rates increase over the length of DNA templates, and that other error processes including PCR misincorporation contribute to the variance of the distribution. These factors contribute to overdispersion. Tests for overdispersion (not shown) indicate the Poisson does not fit the data. This is consistent with the observation that variances exceed means at all loci tested (Table 2). In addition, the Akaike Information Criterion (AIC) and Bayesian Information Criterion (BIC) values consistently show that the negative binomial fit is better than the Poisson fit (Table 2).

**Figure 1.**
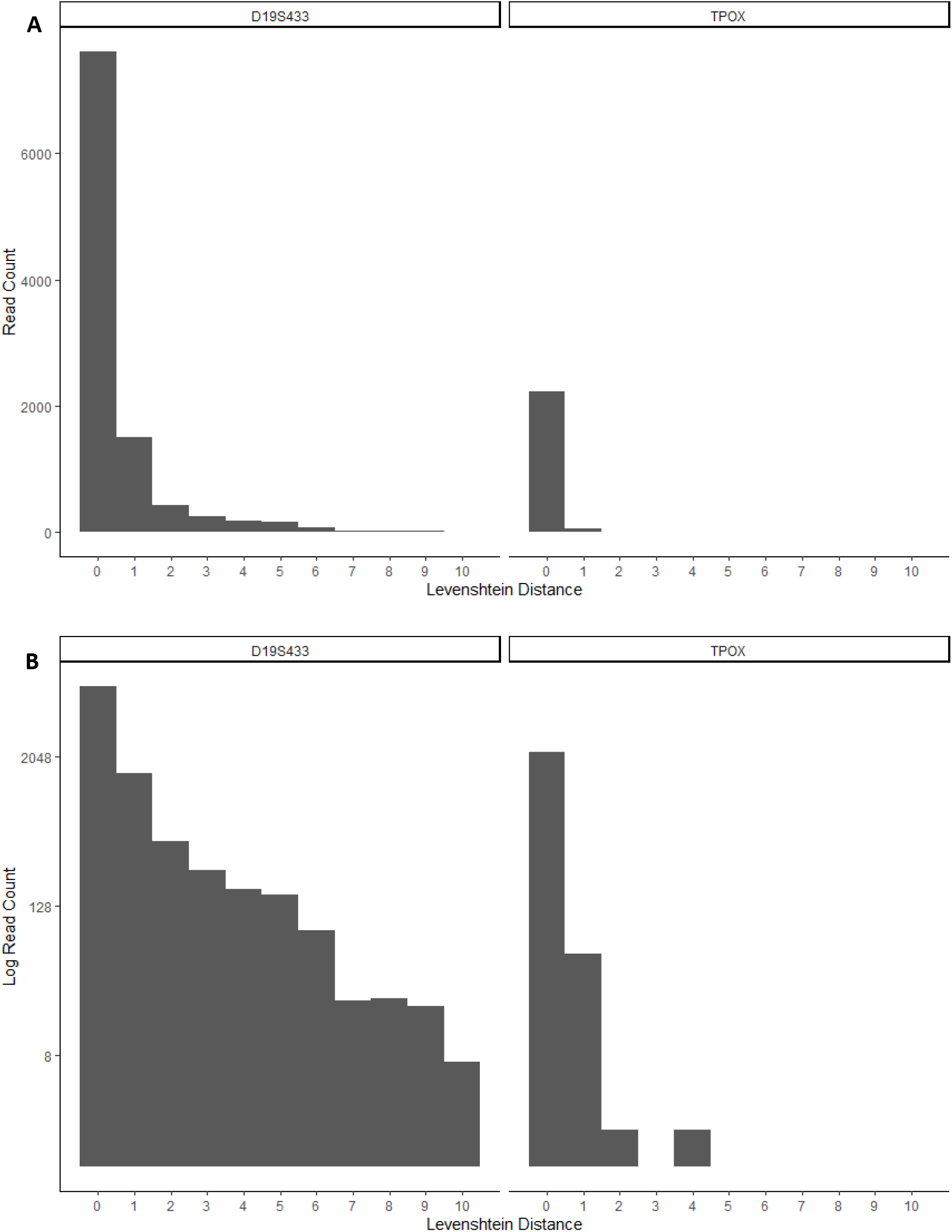
Histograms of read counts from selected loci after filtering stutter. Data was truncated to Levenshtein distances ≤ 10. A) Raw read counts. B) Read counts on log scale to aid visualization of low counts.

**Figure 2.**
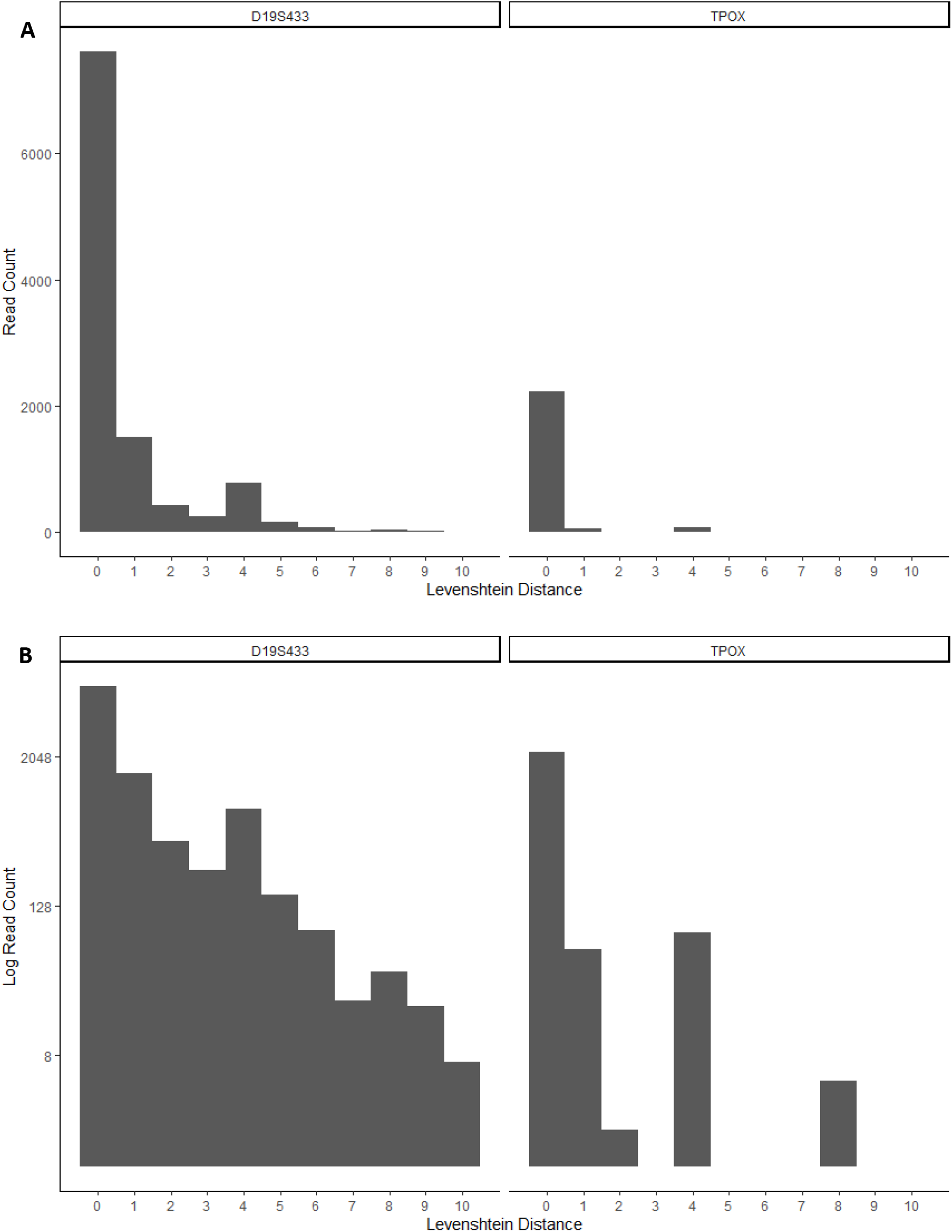
Histograms of read counts from selected loci with stutter included. Data were truncated to Levenshtein distances ≤ 10. A) Raw read counts. B) Read counts on log scale to aid visualization of low counts.

**Table 2.** Read count statistics for 16 loci that were homozygous in the data set.

| Locus | Reads (N) | Mean | Variance | Standard Deviation | Poisson |  | Negative Binomial |  |
| --- | --- | --- | --- | --- | --- | --- | --- | --- |
|  |  |  |  |  | AIC | BIC | AIC | BIC |
| TPOX | 2,296 | 0.02918 | 0.04052 | 0.2013 | 655 | 661 | 606 | 617 |
| D2S441 | 5,809 | 0.06335 | 0.1110 | 0.3332 | 2,918 | 2,925 | 2,699 | 2,713 |
| D3S1358 | 10,822 | 0.1351 | 0.3811 | 0.6173 | 10,088 | 10,095 | 8,092 | 8,106 |
| D4S2408 | 4,881 | 0.03176 | 0.04878 | 0.2209 | 1,430 | 1,436 | 1,334 | 1,347 |
| CSF1PO | 1,170 | 0.06838 | 0.1782 | 0.4221 | 653 | 658 | 531 | 541 |
| D5S818 | 1,062 | 0.06309 | 0.1213 | 0.3483 | 603 | 608 | 493 | 503 |
| D6S1043 | 15,335 | 0.1111 | 0.3090 | 0.5559 | 12,355 | 12,363 | 9,985 | 10,001 |
| D7S820 | 3,747 | 0.1703 | 0.4354 | 0.6598 | 4,074 | 4,080 | 3,347 | 3,359 |
| D9S1122 | 8,670 | 0.03956 | 0.09382 | 0.3063 | 3,134 | 3,141 | 2,605 | 2,619 |
| D10S1248 | 6,407 | 0.1313 | 0.3294 | 0.5739 | 5,741 | 5,748 | 4,824 | 4,837 |
| TH01 | 24,670 | 0.07937 | 0.1824 | 0.4271 | 15,112 | 15,120 | 12,967 | 12,983 |
| D13S317 | 4,185 | 0.1101 | 0.2065 | 0.4544 | 3,185 | 3,192 | 2,906 | 2,918 |
| D19S433 | 10,271 | 0.5516 | 1.643 | 1.282 | 24,843 | 24,850 | 19,489 | 19,504 |
| D20S482 | 18,769 | 0.09414 | 0.2498 | 0.4998 | 13,340 | 13,348 | 10,760 | 10,776 |
| PentaD | 2,292 | 0.2133 | 0.3834 | 0.6192 | 2,731 | 2,737 | 2,567 | 2,578 |
| D22S1045 | 5,501 | 0.1736 | 0.4270 | 0.6534 | 655 | 661 | 606 | 618 |
| **TOTAL** | **125,887** |  |  |  |  |  |  |  |
| **Average** |  | **0.1291** | **0.3213** | **0.5668** |  |  |  |  |

### 3.2. Accounting for Elevated Frequencies of Reads with Levenshtein Distances Equal to The Length of Stutter Motifs After Filtering Stutter

Even when stutter artifacts have been filtered from the data, these artifacts can still contribute to overdispersion in a cryptic way. Frequencies of reads with Levenshtein distances in multiples of tandem repeat motifs are slightly elevated relative to the overall pattern. For example, counts of reads with Levenshtein distances equal to four and eight tend to be slightly elevated for loci with tetramer repeat motifs (Figure 1). This phenomenon is a consequence of defining stutter as a pure motif shift; combined with how Levenshtein distances are calculated. In PCR-MPS, stutter artifacts formed during the PCR step are subject to sequencing error during the sequencing step. If a point sequencing error occurs in a stutter artifact, then a doubly mutated read results where the total number of nucleotide changes is five (assuming a tetramer repeat motif). If the sequencing error occurred in the same repeat motif that was increased or decreased in number due to stutter, then the minimum number of edit operations required to convert the doubly mutated sequence to the correct parent allele sequence may be only four. In the case of N+1 stutter the number of tandem repeats increased by one. When a point error occurs somewhere in the tandem repeat that increased by one repeat, then only the repeat motif containing the sequencing point error need be deleted for an edit distance equal to four. In the case of N−1 stutter, the structure of the repeat motif may be such that the conversion of the doubly mutated sequence to the parent allele sequence can also be solved using only four edits (Appendix C). Elevated frequencies can also be observed for reads with Levenshtein distances of 5 and 9 (assuming a tetramer repeat motif). These elevated frequencies can be attributed to the presence of sequencing errors outside the STR repeat motif of stutter artifacts. In this case, the stutter error necessitates four (or eight) edits, and a point sequencing error necessitates an additional edit. Mechanisms other than those described here may also be involved.

### 3.3. Levenshtein Distance as an Estimator of Error Rates

Locus and profile error rates can be estimated from Levenshtein distances. The mean Levenshtein distances for the sixteen loci studied here ranged from 0.58 errors per 1,000 nucleotides (D9S1122) to 3.73 errors per 1000 nucleotides (D19S433, Table 3). Error rates for most loci were less than 1 per 1,000 nucleotides, which is consistent with published error rates for Illumina sequencing [31]. However, different loci exhibit different error rates, and these differences can be quantified using Levenshtein distances.

**Table 3.** Inference of error rate using Levenshtein distance statistics.

| Locus | Sample | Allele | Allele Length | Mean Levenshtein Distance | Inferred Error Rate |  |
| --- | --- | --- | --- | --- | --- | --- |
|  |  |  |  |  | Per nt | Per 1000 nt |
| TPOX | 2800M | 11 UN | 49 | 0.02918 | 5.96x10<sup>-4</sup> | 0.596 |
| D2S441 | 2391d_A | 11 RC | 96 | 0.06335 | 6.60x10<sup>-4</sup> | 0.660 |
| D3S1358 | 2391d_A | 17 ZU | 134 | 0.1351 | 1.01x10<sup>-3</sup> | 1.01 |
| D4S2408 | 2391d_A | 9 ZI | 50 | 0.03176 | 6.35x10<sup>-4</sup> | 0.635 |
| CSF1PO | 2800M | 12 PD | 68 | 0.06838 | 1.01x10<sup>-3</sup> | 1.01 |
| D5S818 | 2800M | 12 RR | 68 | 0.06309 | 9.28x10<sup>-4</sup> | 0.928 |
| D6D1043 | 2391d_E | 11 BZ | 129 | 0.1111 | 8.61x10<sup>-4</sup> | 0.861 |
| D7S820 | 2391d_B | 10 TO | 95 | 0.1703 | 1.79x10<sup>-3</sup> | 1.79 |
| D9S1122 | 2391d_E | 11 KZ | 68 | 0.03956 | 5.82x10<sup>-4</sup> | 0.582 |
| D10S1248 | 2391d_E | 14 TJ | 97 | 0.1313 | 1.35x10<sup>-3</sup> | 1.35 |
| TH01 | 2391d_B | 7 CS | 72 | 0.07937 | 1.10x10<sup>-3</sup> | 1.10 |
| D13S317 | 2391d_B | 11 AQ | 122 | 0.1101 | 9.02x10<sup>-4</sup> | 0.902 |
| D19S433 | 2391d_E | 14 WU | 148 | 0.5516 | 3.73x10<sup>-3</sup> | 3.73 |
| D20S482 | 2391d_E | 15 QX | 102 | 0.09414 | 9.23x10<sup>-4</sup> | 0.923 |
| PentaD | 2391d_E | 14 WB | 228 | 0.2133 | 9.36x10<sup>-4</sup> | 0.936 |
| D22S1045 | 2800M | 16 AW | 174 | 0.1736 | 9.98x10<sup>-4</sup> | 0.998 |
| **Overall Average** |  |  |  | **0.1291** | **1.13x10<sup>-3</sup>** | **1.13** |

### 3.4. PCR-MPS Method Accuracy and Precision

PCR-MPS method accuracy can be expressed as the mean Levenshtein distance of reads relative to zero. The mean Levenshtein distance is an expression of the degree of conformity of experimentally determined sequences to the expected sequences. In the data presented here, the overall method accuracy is 0.1291 distance units. The standard deviation of Levenshtein distance values is an expression of the degree of mutual agreement among a series of individual sequence determinations. In the data presented here, the overall method precision is the overall standard deviation of 0.5668.

### 3.5. Comparison to Other Measures of Accuracy and Precision

Currently published methods for expressing accuracy and precision in PCR-MPS methods involve either allele call rates [15] or standard deviations of allele read count intensities [32]. These measures are indicators of PCR efficiency and variability but are less informative for whole method accuracy and precision. In many cases where inhibition or low template is not involved, percent call rate can be an insensitive measure (Table 4, Figure 3). Levenshtein distance is more closely related to total method error and is sensitive to differences in method error rates. In this study, we demonstrated the utility of Levenshtein distance-based accuracy and precision by discriminating performances at different loci within a single forensic kit. However, this same method can be employed in the comparison of kits, sequencers, technicians, laboratories, etc.

**Table 4.** Comparison of call rates and standard deviations of read count intensities with Levenshtein distance statistics.

| Locus | Call Rate (%) | Read Count Intensity |  |  | Levenshtein Distance |  |
| --- | --- | --- | --- | --- | --- | --- |
|  |  | Mean | Standard Deviation | Standard Error | Mean | Standard Deviation |
| TPOX | 100 | 722 | 149 | 60.8 | 0.02918 | 0.04052 |
| D2S441 | 100 | 1,884 | 421 | 172.0 | 0.06335 | 0.1110 |
| D3S1358 | 100 | 3,040 | 817 | 333.4 | 0.1351 | 0.3811 |
| D4S2408 | 100 | 1,561 | 507 | 206.8 | 0.03176 | 0.04878 |
| CSF1PO | 100 | 352 | 104 | 42.3 | 0.06838 | 0.1782 |
| D5S818 | 100 | 426 | 146 | 59.5 | 0.06309 | 0.1213 |
| D6D1043 | 100 | 4,745 | 621 | 358.7 | 0.1111 | 0.3090 |
| D7S820 | 100 | 1,204 | 182 | 74.5 | 0.1703 | 0.4354 |
| D9S1122 | 100 | 2,809 | 92 | 53.1 | 0.03956 | 0.09382 |
| D10S1248 | 100 | 1,948 | 142 | 82 | 0.1313 | 0.3294 |
| TH01 | 100 | 7,571 | 1,378 | 562.4 | 0.07937 | 0.1824 |
| D13S317 | 100 | 1,482 | 559 | 228.3 | 0.1101 | 0.2065 |
| D19S433 | 100 | 2,537 | 892 | 515.0 | 0.5516 | 1.643 |
| D20S482 | 100 | 5,876 | 687 | 396.8 | 0.09414 | 0.2498 |
| PentaD | 100 | 639 | 315 | 181.8 | 0.2133 | 0.3834 |
| D22S1045 | 100 | 1,942 | 420 | 171.5 | 0.1736 | 0.4270 |
| **Average** | **100** | **2,421** | **464** | **218.7** | **0.1291** | **0.3213** |

**Figure 3.**
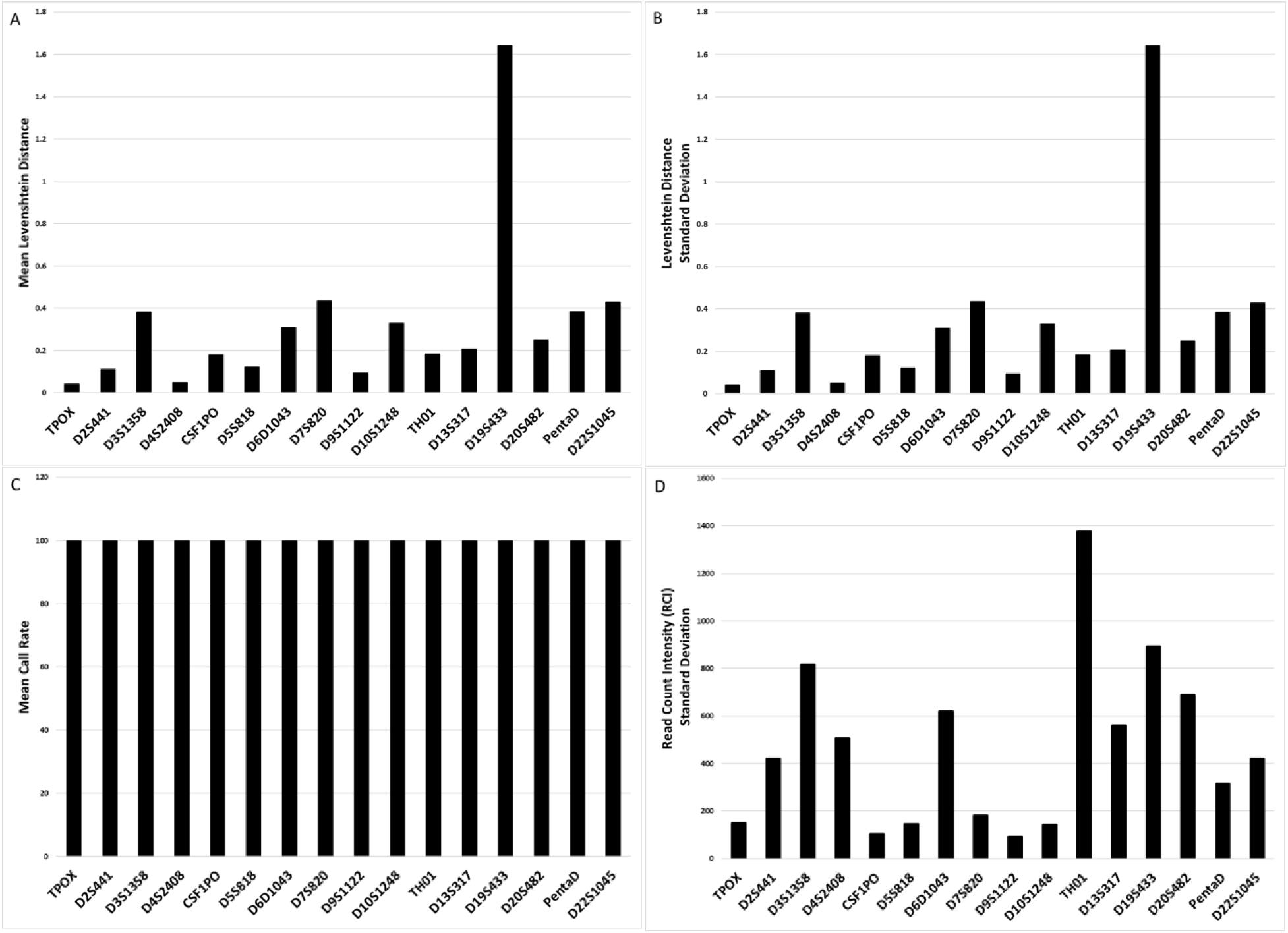
Alternative measures of accuracy and precision in PCR-MPS methods. Each plot represents data pooled across triplicate analyses of DNA standards. A) The mean Levenshtein distance is a measure of method accuracy where larger values indicate lower method accuracy. B) The standard deviation of Levenshtein distance is a measure of method precision where larger values correspond to lower precision. C) Call rates. D) Standard deviations of read count intensities.

## 4. Conclusions

Levenshtein distance is a metric that can be used to calculate accuracy and precision of forensic PCR-MPS methods in compliance with SWGDAM and ENSFI guidelines. The Levenshtein distance has an intuitive interpretation as the edit distance between experimentally determined reads and expected or reference sequences.

## Acknowledgements

Supported by Army SBIR W911NF-20-P-0068

## Appendix A

Definitions for mean and variance imply the need for a metric that can be used to quantitatively calculate accuracy and precision. A measure that satisfies the axioms of a metric is essential for calculating accuracy and precision in a way analogous to the approach used for PCR-CE methods, including calculation of standard deviations. Metrics are distance functions that must satisfy four axioms [25] p.147. Let *d*(*x*, *y*) be a function measuring the distance between two elements ***x*** and ***y*** in a set of elements S.

1. The distance between two elements in the metric space is non-negative: *d*(*x*, *y*) ≥ 0
2. The distance between two entities is zero if and only if the two entities are the same: *d*(*x*, *x*) = 0
3. The order of comparison does not matter: *d*(*x*, *y*) = *d*(*y*, *x*)
4. The distance between two elements is less than or equal to the sum of the distances obtained when an intermediate element is inserted. In other words, the triangle inequality is satisfied: *d*(*x*, *z*) ≤ *d*(*x*, *y*) + *d*(*y*, *z*)

These properties hold for common metrics such as meters or inches in metric spaces of ordinary experience. These properties also hold when the metric *d*(*x*, *y*) is Levenshtein distance and the elements in the metric space are DNA sequence strings [24].

## Appendix B

**Figure B-1.**
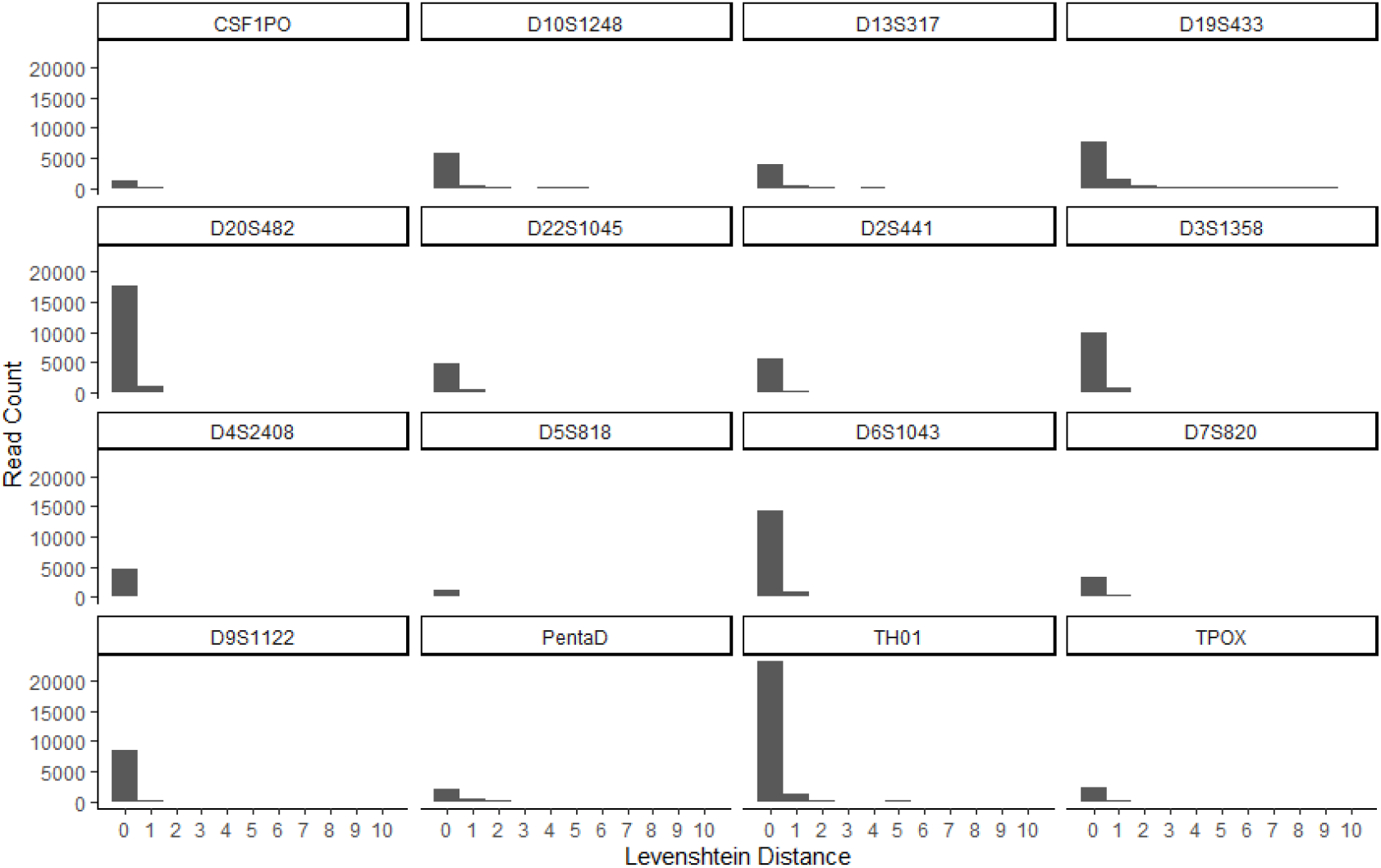
Histograms for sixteen loci in normal scale.

**Figure B-2.**
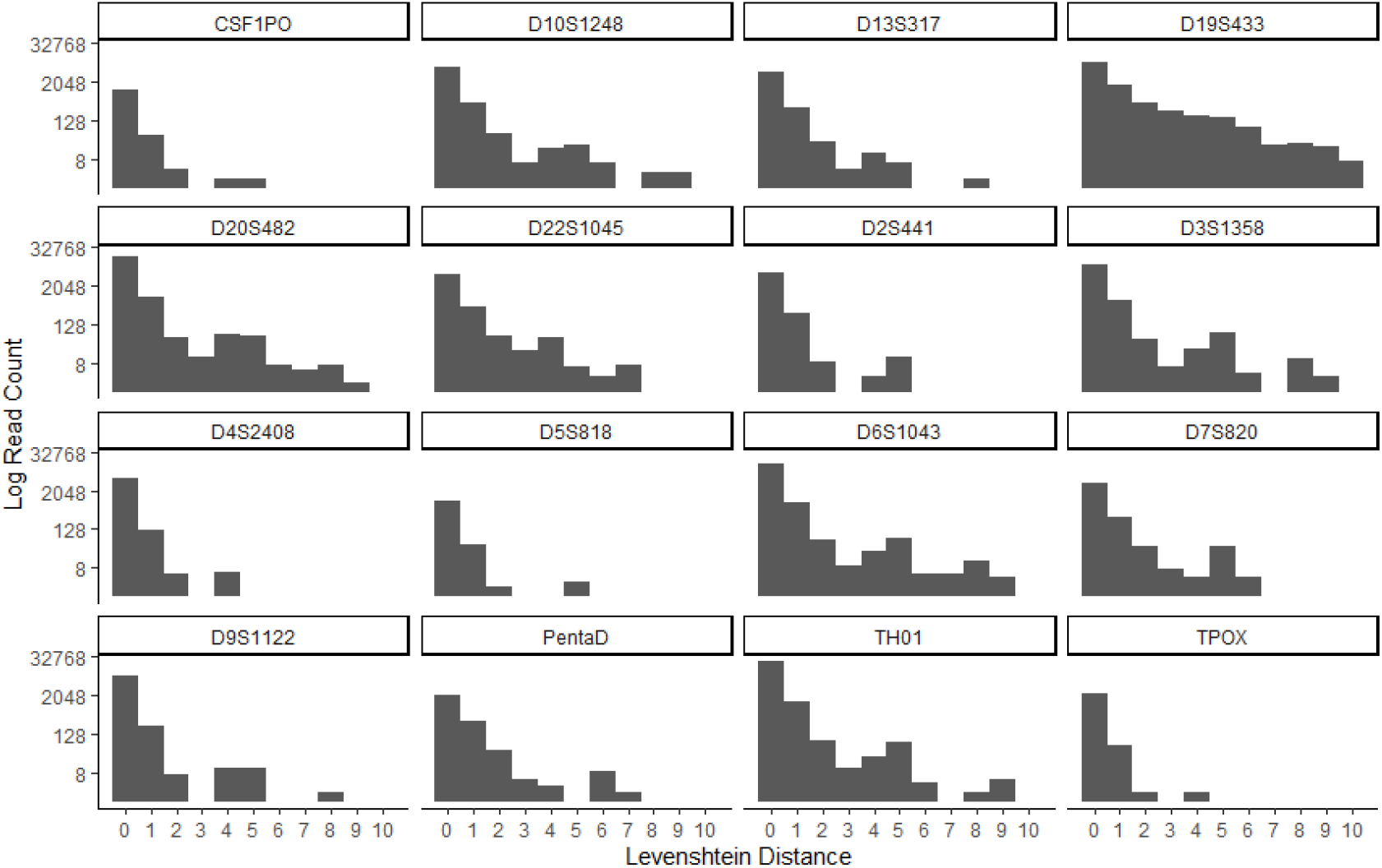
Histograms for sixteen loci in log scale.

## Appendix C Explanation of Elevated Frequencies of Levenshtein Distances of Four and Five at Loci for Tetrameric Tandem Repeat Motifs

**Figure C-1.**
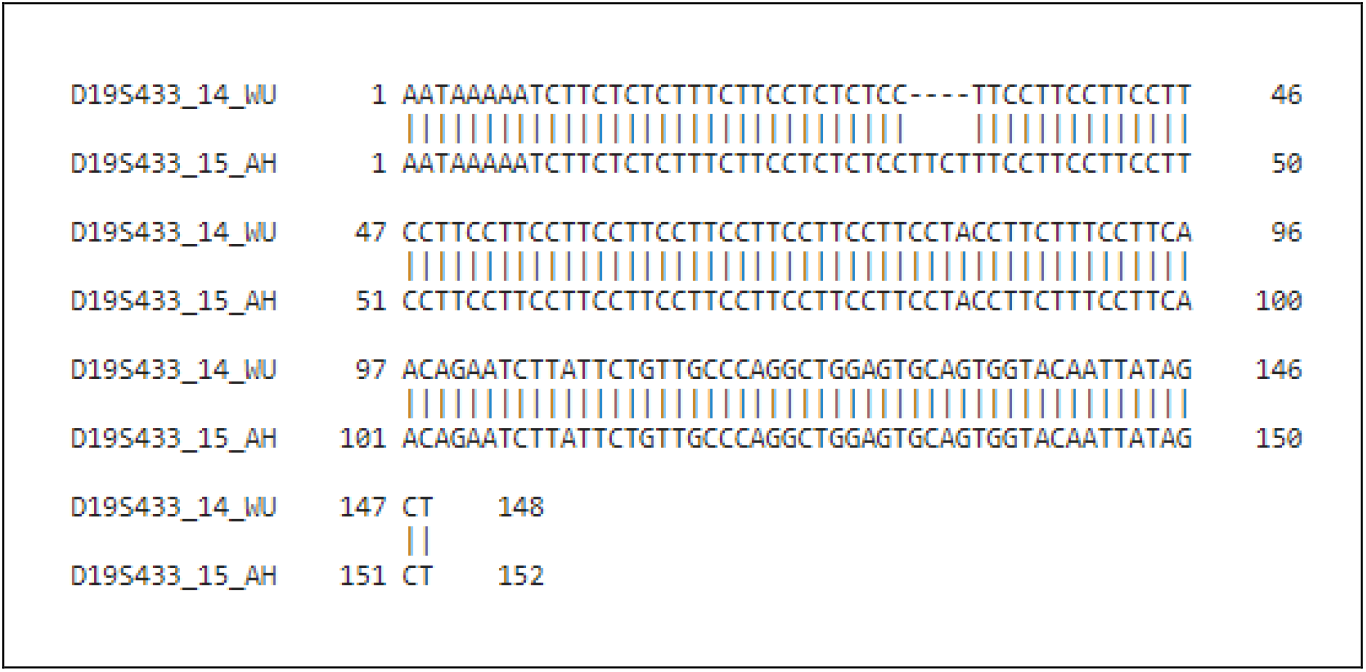
Alignment of the D19S433 14 WU and 15 AH sequences. The 15 AH sequence is an N+1 stutter artifact of the 14 WU allele sequence with the addition of a sequencing error inside the repeat motif of the stutter artifact. Alignment can be solved using only four edit operations. The bracketed STR sequences are [CCTT]_12_[CCTA]_1_[CCTT]_1_[CTTT]_1_[CCTT]_1_ and [CCTT]_1_[CTTT]_1_[CCTT]_11_[CCTT]_1_[CCTT]_1_[CTTT]_1_[CCTT]_1_ respectively. Alignment was performed using EMBOS Needle with default parameter values.

**Figure C-2.**
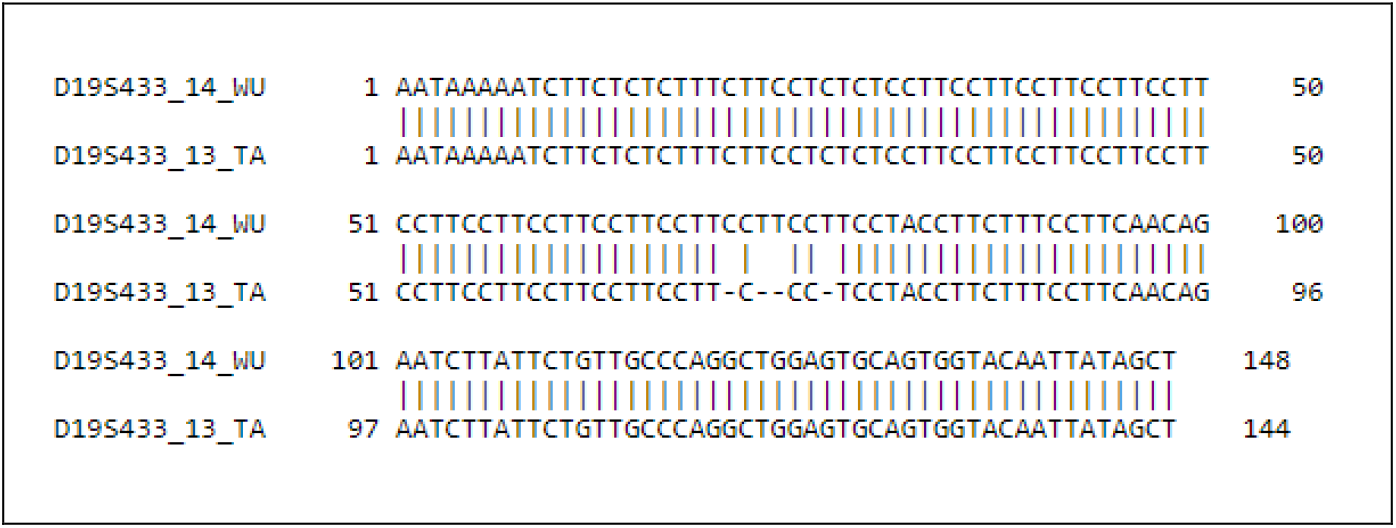
Alignment of the D19S433 14 WU and 13 TA sequences. The 13 TA sequence is an N-1 stutter artifact of the 14 WU allele sequence with the addition of a sequencing error inside the repeat motif of the stutter artifact. Alignment can be solved using only four edit operations. The bracketed STR sequences are [CCTT]_12_[CCTA]_1_[CCTT]_1_[CTTT]_1_[CCTT]_1_ and [CCTT]_10_CCCT[CCTA]_1_[CCTT]_1_[CTTT]_1_[CCTT]_1_ respectively. Alignment was performed using EMBOS Needle, with gap open = 5.0 and gap extend = 0.0005.

**Figure C-3.**
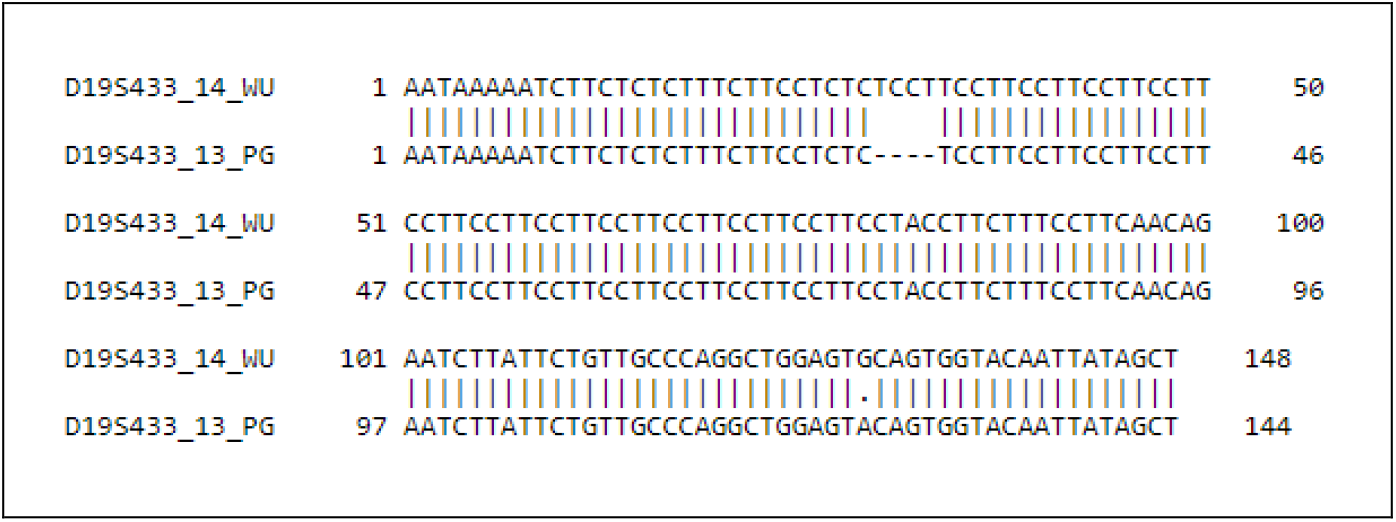
Alignment of the D19S433 14 WU and 13 PG sequences. The 13 PG sequence is an N-1 stutter artifact of the 14 WU allele sequence with the addition of a sequencing error outside the repeat motif of the stutter artifact. Alignment can be solved using five edit operations. The bracketed STR sequences are [CCTT]_12_[CCTA]_1_[CCTT]_1_[CTTT]_1_[CCTT]_1_ and [CCTT]_11_[CCTA]_1_[CCTT]_1_[CTTT]_1_[CCTT]_1_ respectively. Alignment was performed using EMBOS Needle, with default parameter values.

**Figure C-4.**
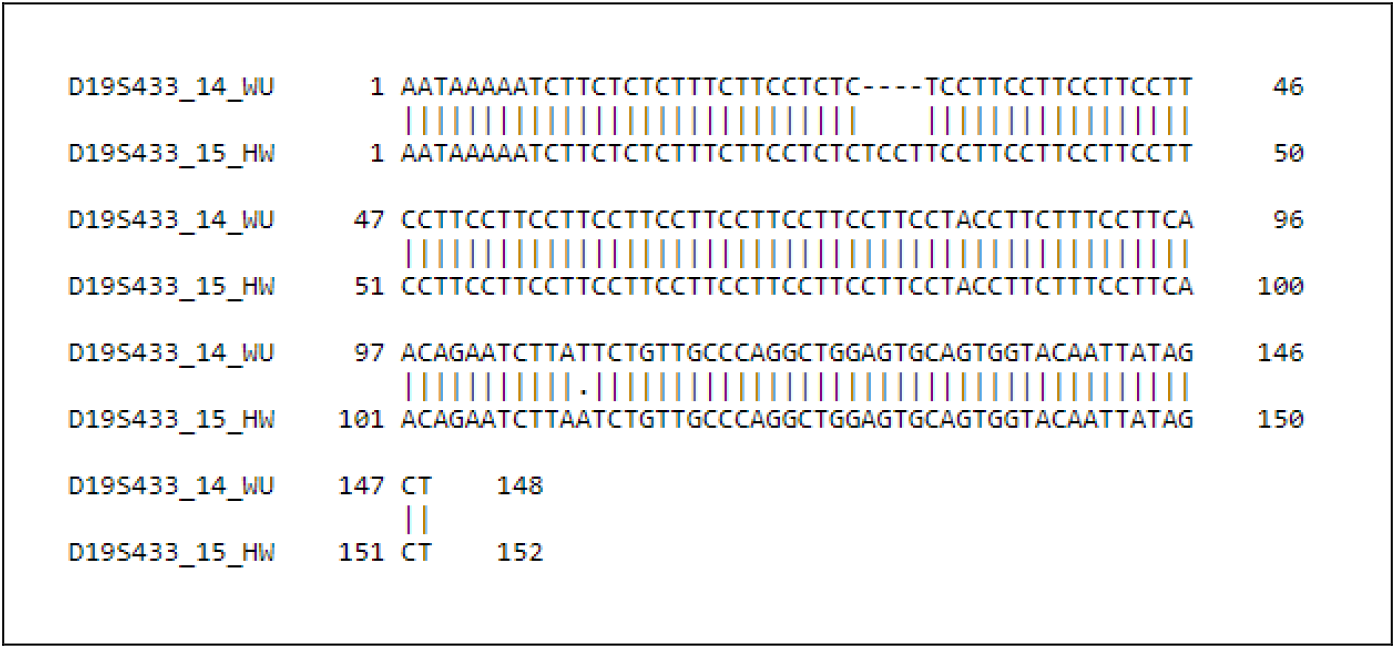
Alignment of the D19S433 14 WU and 15HW sequences. The 15 HW sequence is an N+1 stutter artifact of the 14 WU allele sequence with the addition of a sequencing error outside the repeat motif of the stutter artifact. Alignment can be solved using five edit operations. The bracketed STR sequences are [CCTT]_12_[CCTA]_1_[CCTT]_1_[CTTT]_1_[CCTT]_1_ and [CCTT]_13_[CCTA]_1_[CTTT]_1_[CCTT]_1_[CCTT]_1_ respectively. Alignment was performed using EMBOS Needle, with default parameter values.

## Footnotes

1 The IUPAC ambiguity code N can appear in sequencer output but reads containing N characters are typically not considered in forensic analysis and will be disregarded here.

2 Our use of the term nucleotide is functionally equivalent to the term base pair commonly used in forensics. The term nucleotide is preferred because the object of most bioinformatic analysis is the sequencer read, which represents just one DNA strand.

## References

[1] S. Bell, Measurement uncertainty in forensic science : a practical guide, CRC Press, Taylor & Francis Group, Boca Raton, 2017.

[2] T. Vosk, A.F. Emery, Forensic metrology : scientific measurement and inference for lawyers, judges and criminalists, CRC Press,Taylor & Francis Group, Boca Raton, 2015.

[3] SWGDAM, Scientific Working Group on DNA Analysis Methods Validation Guidelines for DNA Analysis Methods, 2016.

[4] SWGDAM, Addendum to “SWGDAM Interpretation Guidelines for Autosomal STR Typing by Forensic DNA Testing Laboratories” to Address Next Generation Sequencing, 2019.

[5] ENFSI, Recommended Minimum Criteria for the Validation of Various Aspects of the DNA Profiling Process, 2010.

[6] D.Y. Wang, C.W. Chang, R.E. Lagace, L.M. Calandro, L.K. Hennessy, Developmental validation of the AmpFlSTR(R) Identifiler(R) Plus PCR Amplification Kit: an established multiplex assay with improved performance, J Forensic Sci 57(2) (2012) 453–65.

[7] R.L. Green, R.E. Lagace, N.J. Oldroyd, L.K. Hennessy, J.J. Mulero, Developmental validation of the AmpFlSTR(R) NGM SElect PCR Amplification Kit: A next-generation STR multiplex with the SE33 locus, Forensic Sci Int Genet 7(1) (2013) 41–51.

[8] T.R. Moretti, A.L. Baumstark, D.A. Defenbaugh, K.M. Keys, A.L. Brown, B. Budowle, Validation of STR typing by capillary electrophoresis, J Forensic Sci 46(3) (2001) 661–76.

[9] R.M. Goor, L. Forman Neall, D. Hoffman, S.T. Sherry, A mathematical approach to the analysis of multiplex DNA profiles, Bull Math Biol 73(8) (2011) 1909–31.

[10] M.G. Ensenberger, K.A. Lenz, L.K. Matthies, G.M. Hadinoto, J.E. Schienman, A.J. Przech, M.W. Morganti, D.T. Renstrom, V.M. Baker, K.M. Gawrys, M. Hoogendoorn, C.R. Steffen, P. Martin, A. Alonso, H.R. Olson, C.J. Sprecher, D.R. Storts, Developmental validation of the PowerPlex((R)) Fusion 6C System, Forensic Sci Int Genet 21 (2016) 134–44.

[11] L.K. Hennessy, N. Mehendale, K. Chear, S. Jovanovich, S. Williams, C. Park, S. Gangano, Developmental validation of the GlobalFiler((R)) express kit, a 24-marker STR assay, on the RapidHIT((R)) System, Forensic Sci Int Genet 13 (2014) 247–58.

[12] I.-I.C.o.B. Nomenclature, Abbreviations and symbols for nucleic acids, polynucleotides, and their constituents, Biochemistry 9(20) (1970) 4022–4027.

[13] K.B. Gettings, L.A. Borsuk, D. Ballard, M. Bodner, B. Budowle, L. Devesse, J. King, W. Parson, C. Phillips, P.M. Vallone, STRSeq: A catalog of sequence diversity at human identification Short Tandem Repeat loci, Forensic Sci Int Genet 31 (2017) 111–117.

[14] K.B. Gettings, L.A. Borsuk, C.R. Steffen, K.M. Kiesler, P.M. Vallone, Sequence-based U.S. population data for 27 autosomal STR loci, Forensic Sci Int Genet 37 (2018) 106–115.

[15] A.C. Jager, M.L. Alvarez, C.P. Davis, E. Guzman, Y. Han, L. Way, P. Walichiewicz, D. Silva, N. Pham, G. Caves, J. Bruand, F. Schlesinger, S.J.K. Pond, J. Varlaro, K.M. Stephens, C.L. Holt, Developmental validation of the MiSeq FGx Forensic Genomics System for Targeted Next Generation Sequencing in Forensic DNA Casework and Database Laboratories, Forensic Sci Int Genet 28 (2017) 52–70.

[16] R.S. Just, L.I. Moreno, J.B. Smerick, J.A. Irwin, Performance and concordance of the ForenSeq system for autosomal and Y chromosome short tandem repeat sequencing of reference-type specimens, Forensic Sci Int Genet 28 (2017) 1–9.

[17] J.D. Churchill, S.E. Schmedes, J.L. King, B. Budowle, Evaluation of the Illumina((R)) Beta Version ForenSeq DNA Signature Prep Kit for use in genetic profiling, Forensic Sci Int Genet 20 (2016) 20–29.

[18] J.D. Churchill, J. Chang, J. Ge, N. Rajagopalan, S.C. Wootton, C.W. Chang, R. Lagace, W. Liao, J.L. King, B. Budowle, Blind study evaluation illustrates utility of the Ion PGM system for use in human identity DNA typing, Croat Med J 56(3) (2015) 218–29.

[19] S. Kocher, P. Muller, B. Berger, M. Bodner, W. Parson, L. Roewer, S. Willuweit, D.N. Consortium, Inter-laboratory validation study of the ForenSeq DNA Signature Prep Kit, Forensic Sci Int Genet 36 (2018) 77–85.

[20] V. Sharma, H.Y. Chow, D. Siegel, E. Wurmbach, Qualitative and quantitative assessment of Illumina's forensic STR and SNP kits on MiSeq FGx, PLoS One 12(11) (2017) e0187932.

[21] A.L. Silvia, N. Shugarts, J. Smith, A preliminary assessment of the ForenSeq FGx System: next generation sequencing of an STR and SNP multiplex, Int J Legal Med 131(1) (2017) 73–86.

[22] D. Gusfield, Algorithms for Strings, Trees, and Sequences, Cambridge University Press, Cambridge, 1997.

[23] V. Levenshtein, Binary codes capable of correcting deletions, insertions, and reversals, Soviet Physics Doklady 10(8) (1965) 707–710.

[24] G. Navarro, A guided tour to approximate string matching, ACM Comput. Surv. 33(1) (2001) 31–88.

[25] S. Axler, Measure, Integration & Real Analysis, Springer, Cham, Switzerland, 2020.

[26] J. Maynard Smith, Natural selection and the concept of a protein space, Nature 225(February 7) (1970) 563–564.

[27] M. Eigen, R. Winkler-Oswatitsch, A. Dress, Statistical geometry in sequence space: a method of quantitative comparative sequence analysis, Proc Natl Acad Sci U S A 85(16) (1988) 5913–7.

[28] A.R. Isenberg, R.O. Allen, K.M. Keys, J.B. Smerick, B. Budowle, B.R. McCord, Analysis of two multiplexed short tandem repeat systems using capillary electrophoresis with multiwavelength fluorescence detection, Electrophoresis 19(1) (1998) 94–100.

[29] K.B. Gettings, D. Ballard, M. Bodner, L.A. Borsuk, J.L. King, W. Parson, C. Phillips, Report from the STRAND Working Group on the 2019 STR sequence nomenclature meeting, Forensic Sci Int Genet 43 (2019) 102165.

[30] B. Young, T. Faris, L. Armogida, A nomenclature for sequence-based forensic DNA analysis, Forensic Sci Int Genet 42 (2019) 14–20.

[31] M. Schirmer, U.Z. Ijaz, R. D’Amore, N. Hall, W.T. Sloan, C. Quince, Insight into biases and sequencing errors for amplicon sequencing with the Illumina MiSeq platform, Nucleic Acids Res 43(6) (2015) e37.

[32] R. Tao, W. Qi, C. Chen, J. Zhang, Z. Yang, W. Song, S. Zhang, C. Li, Pilot study for forensic evaluations of the Precision ID GlobalFiler NGS STR Panel v2 with the Ion S5 system, Forensic Sci Int Genet 43 (2019) 102147.

